# Characterization of Recombinant Human Lactoferrin Expressed in *Komagataella Phaffii*

**DOI:** 10.1101/2024.01.09.574900

**Authors:** Xiaoning Lu, Chad Cummings, Udodili A. Osuala, Neela H. Yennawar, Kevin E. W. Namitz, Brittney Hellner, Pamela B. Besada-Lombana, Ross D. Peterson, Anthony J. Clark

## Abstract

We performed a thorough analysis and characterization of multiple batches of Helaina recombinant human lactoferrin (rhLF, Effera™) expressed at an industrial scale in a yeast system. Bottom-up LC-MS/MS-based proteomics analysis detected the full sequence of Helaina rhLF protein and confirmed that its amino acid sequence is identical to that of native human LF (Uniprot i.d. P02788). Helaina rhLF had a protein purity of 98% or higher as determined by three orthogonal methods; reversed-phase HPLC, SDS-PAGE, and LC-MS proteomics analysis. N-linked glycans were detected at three known glycosylation sites, namely, Asparagines-156, -497, and -642. The identified N-glycans of Helaina rhLF were predominantly oligomannose structures with five to nine mannoses (M5-M9), which we also report to be present in both the native human and bovine LF. human milk LF (hmLF) possessed lower levels of oligomannose structures and were mainly M5 and M6. Helaina rhLF protein secondary structure was nearly identical to that of hmLF, as revealed by microfluidic modulation spectroscopy. Results of small-angle X-ray scattering (SAXS) and analytical ultracentrifugation analyses confirmed that, like hmLF, Helaina rhLF displayed well-folded globular structures in solution. Reconstructed solvent envelopes of Helaina rhLF, obtained through the SAXS analysis, demonstrated a remarkable fit with the reported crystalline structure of iron-bound native hmLF. Differential scanning calorimetry investigations into the thermal stability of Helaina rhLF revealed two distinct denaturation temperatures at 68.7±0.9 °C and 91.9±0.5 °C, consistently mirroring denaturation temperatures observed for apo-and holo-hmLF. Overall, the characterization analysis results affirmed that Helaina rhLF was of high purity and exhibited globular structures closely akin to that of hmLF.

## INTRODUCTION

Human lactoferrin (hLF) is an iron-binding glycoprotein found in high concentrations in human milk and to a lesser degree in tears, saliva, intestinal secretions, and neutrophils [1]. Human milk LF (hmLF) is a bioactive protein that nourishes newborns and supports optimal growth and development [2, 3]. Lactoferrin (LF) also supports immune health at every life stage [4, 5]. It is the second most abundant glycoprotein in human milk, constituting an estimated 15% of the total whey protein of human milk [6]. Human milk LF is a single polypeptide chain containing 691 amino acid residues with a molecular weight (MW) of about 80 kDa, and its secondary structure comprises α-helices, β-pleated sheets, and turns [7]. Its structure consists of two folded lobes connected by a three-turn α-helix, with each lobe having the capacity to bind one ferric ion (Fe^3+^) [7, 8], a characteristic that may assist iron absorption in the small intestine of infants fed human milk [1, 9].

N-glycosylation, a common yet complex post-translational modification, is another characteristic of hmLF with important structural and functional roles [10, 11]. The N-glycans are linked to the asparagine (Asn) residue of hmLF, which, in turn, affects its structure and potential biological function [11, 12]. In nature, N-glycan structures of LF vary across cell types (e.g., human milk-derived vs neutrophil-derived) and species (e.g., human vs bovine), from mother to mother, and throughout the lactation period [13, 14]. For example, hmLF contains primarily complex N-glycans that are frequently sialylated and fucosylated and low abundance of oligomannose [15], while bovine lactoferrin (bLF) has a different glycan distribution that includes abundant oligomannose, complex and hybrid glycans [16]. In summary, hmLF contains more complex N-glycans than bLF, which contains all three glycan types (Fig. 1A) [15, 17]. In addition, glycosylation in yeast generally results in hyper-mannosylated glycoforms [18], a characteristic not found in mammalian systems (Fig. 1B).

**Fig. 1.**
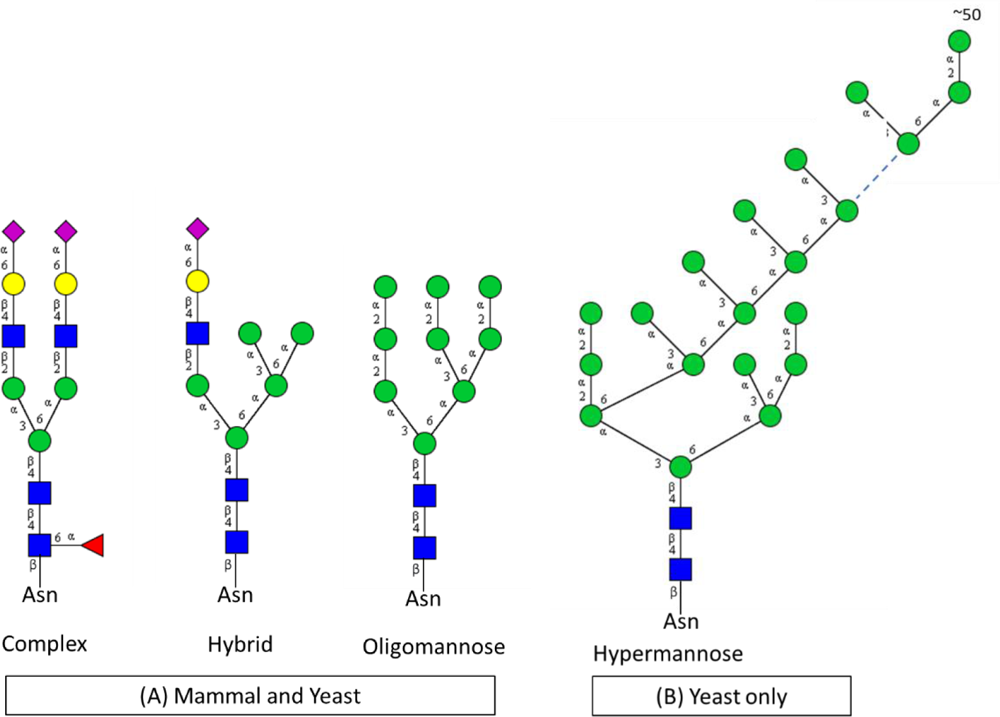
The three types of N-glycans that are present in the glycoproteins of both mammals and yeast, namely complex, hybrid, and oligomannose structures (A). The N-glycans share a common core Man3GlcNAc2 which is linked to the Asparagine (Asn) residue of the glycoproteins having Asparagine (Asn)-X-Serine (Ser)/Threonine (Thr) sequons. Unlike mammals, the yeast tends to synthesize hypermannose N-Glycans (B). The colored shapes represent monosaccharide residues: 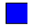 N-acetylglucosamine (GlcNAc); 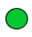 mannose (Man); 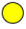 galactose (Gal), 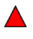 fucose (Fuc); and 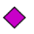 N-acetylneuraminic acid (NeuAc).

Data from preclinical and clinical studies have shown the effects of supplementing the diet with LF on markers of iron status, highlighting the potential benefits of hmLF, bLF, and recombinant hLF (rhLF) as a food ingredient [19–23]. A preclinical study demonstrated that supplementation with rhLF from transgenic cows improved iron status markers (e.g., red blood cells, hemoglobin, serum iron, serum ferritin) [19]. Clinical studies have shown that extended dietary supplementation with LF increased serum iron, red blood cell number, and serum ferritin [20, 21], findings that have been confirmed in a recently published systematic review and meta-analysis of 11 intervention studies [22].

In light of these declared benefits, there is a desire to use recombinant genetic technology to develop sustainable methods for the industrial scale production of rhLF. Indeed, a variety of recombinant systems have been used to express rhLF efficiently, including transgenic animals (e.g., cows), plants (e.g., rice), filamentous fungi (e.g., *Aspergillus niger* var*. awamori*), and yeast (e.g., *Saccharomyces cerevisiae*, *Komagataella phaffii* (*K. phaffii*, formerly classified as *Pichia pastoris*) [24–31]. For example, rhLF expressed from transgenic animal systems has a molecular weight, protein structure, and function similar to hmLF, yet their glycan structures differ [15]. Both hmLF and rhLF from transgenic cows contain sialylated and fucosylated glycans, whereas oligomannose glycans were reported with high abundance in the rhLF from transgenic cows and low abundance in hmLF [11, 15].

*K. phaffii* is a eukaryotic, methylotrophic yeast commonly used to express recombinant proteins efficiently, including animal-derived and human-derived recombinant LF [32–35]. Unlike most recombinant systems, *K. phaffii* can be modified to produce rhLF with glycan structures similar to hmLF. Also distinct from other host expression systems, *K. phaffii* can carry out high-density fermentation, which is a cost-effective way to generate high yields of recombinant proteins [33]. *K. phaffii* has been used as a production organism for numerous food ingredients that have received “no questions” letters from the U.S. Food and Drug Administration in the generally recognized as safe (GRAS) notification program[36–38]. Helaina has successfully used a proprietary technology to produce a modified strain of *K. phaffii* capable of expressing a form of rhLF substantively equivalent to hmLF at an industrial scale for the first time. Using multiple analytical methods, this publication comprehensively characterizes the identity, quality, glycosylation, structure, and iron saturation of Helaina rhLF.

## MATERIALS and METHODS

### Materials

Samples of hmLF from human milk were supplied by MilliporeSigma (St. Louis, MO, USA) and Abcam (Waltham, Boston, USA); bovine LF (bLF) was supplied by MilliporeSigma or The Lactoferrin Company (Byron Bay, Australia). Additionally, two hmLF samples were isolated and purified in-house from two different sources of human milk according to an established protocol [39, 40]. Other reagents were purchased from MilliporeSigma or Thermo Fisher Scientific (Waltham, MA, USA) and used as is.

## Methods

### Production of Helaina rhLF

Multiple batches of Helaina rhLF (Effera™) were produced from a *K. phaffii* fermentation process at an industrial scale. The fermentation process consisted of batch, fed-batch, and induction phases. At the end of fermentation, cells were separated from the protein-containing broth. Recombinant hLF was then isolated by microfiltration, ultrafiltration/diafiltration and ion exchange chromatography. The resulting LF-rich solution was finally spray-dried to a powder. Samples are available upon request.

### SDS-PAGE and Western Blot

SDS-PAGE separation was performed for 42 min at 200V under denaturing and reducing conditions with NuPAGE 4-12% Bis-Tris 1.0 mm Gel (Thermo Fisher Scientific) [41]. Gels were imaged after colorimetric staining with Imperial™ Protein Stain (Coomassie dye R-250). Following SDS-PAGE, Proteins separated in the gel were transferred to a PVDF membrane for LF-specific detection via immunofluorescent staining. Following protein transfer, membranes were incubated with rabbit anti-lactoferrin monoclonal antibody (Millipore Sigma), then with Alexa Fluor™ 488 goat anti-rabbit IgG (Thermo Fisher Scientific). Protein samples were visualized with fluorescent imaging (excitation: 455-485 nm; emission: 508-557 nm).

### Reversed-phase HPLC determination of LF

Protein samples were analyzed using a 1290 UHPLC with a diode-array detector and an AdvanceBio RP-mAb C4 column (2.1 x 75 mm, 3.5 µm, Agilent, Santa Clara, CA). Mobile phases were (A) water with 0.1% trifluoroacetic acid and, (B) acetonitrile (ACN) with 0.09% trifluoroacetic acid. The gradient increased from 25% (B) to 35% (B) in 2 min, then to 45% (B) in 5 min and 55% (B) in 0.5 min, back to 25% (B) in 0.5 min, and was kept at 25% (B) for 2 min to re-equilibrate the column. The flow rate was 0.5 mL/min; the column temperature was set at 70 °C; and detection was set at 214 nm and 280 nm. The calibration standard was hmLF protein diluted in 50 mM Tris (pH 8.0). Peak area, concentration, peak retention time, peak area percentage, and calibration curve parameters were analyzed using ChemStation software.

### Liquid chromatography and tandem mass spectrometry (LC-MS/MS)

#### A. Protein identification of rhLF with full protein sequence coverage

LF was solubilized in 1 mL Tris HCl buffer (pH 8.0). Samples were diluted 3 × 10 μg in 25 mM ammonium carbonate and incubated with 10 mM dithiothreitol at 60 °C, followed by 50 mM iodoacetamide at room temperature. Each aliquot of the samples was digested with trypsin, chymotrypsin, and elastase enzymes (Promega, Madison, WI, USA), separately, at 37 °C for 18 hr, quenched with formic acid, and desalted using solid-phase extraction.

Protein digests were subjected to analysis by nano-flow LC-MS/MS on an Orbitrap Fusion Lumos Tribrid mass spectrometer (Thermo Fisher Scientific) coupled with a Waters^TM^ M-class HPLC system (Milford, MS, USA). Peptides were loaded onto a trapping column and eluted over a 75 μm analytical column at 350 nL/min; both columns were packed with Luna C18 resin (Phenomenex, Torrance, CA, USA). A 30-min gradient was applied. The mass spectrometer was operated in data-dependent mode, with MS and MS/MS performed at resolutions of 60,000 and 15,000 full width half maximum, respectively. Advanced peak determination was enabled. The instrument was run using a 3-second cycle for MS and MS/MS.

The acquired MS data was processed using a local copy of the Byonic^TM^ MS/MS search engine (Protein Metrics, Cupertino, CA, USA) with the following parameters: trypsin or none was set as the enzyme; a UniProt library containing sequencing information for *K. phaffii* and hLF was generated for protein sequencing; carbamidomethylation of cysteine (C) was specified as the fixed modification; and oxidation of methionine (M), acetylation of protein N-terminals, deamidation of asparagine (N) and glutamine (Q), phosphorylation of serine (S), threonine (T), and tyrosine (Y). Additional identification parameters included monoisotopic mass values with a peptide mass tolerance of 10 ppm, fragment mass tolerance of 20 ppm, and max missed cleavages of two. Byonic mzid files were parsed into the Scaffold software version 5.3.0 to validate, filter, and create a nonredundant list for each sample. Data was filtered using criteria that included a minimum protein value of 99%, a minimum peptide value of 50% (Prophet scores), and a minimum of two unique peptides per protein.

#### B. Site-specific glycosylation

Similar LC-MS/MS-based proteomics analyses were performed to determine site-specific glycosylation of rhLF proteins using previously established methods with minimal modification [42–44]. Briefly, an aliquot of 20 µg of rhLF proteins was reduced and alkylated and then digested with trypsin, followed by a cleanup by solid phase extraction using SOLA HRP 10 mg × 1 ml cartridges. Resulting protein digests were analyzed using an Orbitrap Fusion Lumos Tribrid mass spectrometer coupled with a Dionex UltiMate 3000 RSLCnano system (Thermo Fisher Scientific). All MS and MS/MS raw spectra from each sample were searched using Byonic^TM^ MS/MS search engine against the *K. phaffii* NCBI database with added targeted protein hLF sequence. The peptide search parameters were as follows: two missed cleavages for full trypsin digestion with fixed carbamidomethyl modification of cysteine and variable modifications, including methionine oxidation and deamidation on asparagine/glutamine residues. Mass tolerance for peptides was set at 10 ppm, and fragments generated by HCD and EThcD at 0.05 Da and 0.6 Da, respectively. The maximum number of common and rare modifications was set at two. The glycan search was performed against a list of 309 mammalian N-linked glycans. Identified peptides were filtered for a maximum 1% false discovery rate.

The glycan occupancy rate of each site was calculated by dividing the total peak areas of the glycopeptides by the total peak areas of all peptides, including the identified glycopeptides and their respective non-glycosylated peptides as follows:

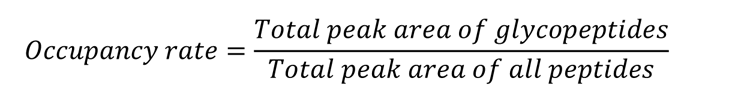

This provided an estimated occupancy rate and assumed the glycopeptides and their respective non-glycosylated peptides produced MS signals with similar efficiency.

#### Released N-glycan analysis by HPLC with fluorescence and mass spectrometry

N-glycans were released, labeled, and isolated from intact protein using the Agilent Gly-X InstantPC kit according to the manufacturer’s instructions (Agilent application note 5994-3482EN). Samples were analyzed on an Agilent 6545XT AdvanceBio LC/Q-TOF instrument equipped with a fluorescence detector (excitation: 285 nm, emission: 345 nm) coupled with an AdvanceBio glycan mapping column (300Å, 2.1 × 150, 1.8 μm). Alternatively, an Agilent 1290 Infinity UHPLC with a fluorescence detector was employed to record just the glycan profiles. Mobile phases were 100 mM ammonium formate, pH 4.5 (A) and ACN (B). Glycan separation was carried out with a gradient from 80% to 60% (A) over 32 min at a constant elution rate of 0.5 mL/min and column temperature of 35 °C. Glycans were identified by matching their observed masses with the theoretical masses of 201 known InstantPC-labeled N-glycans with a mass tolerance of 10 ppm. The glycan identification was further verified by the HPLC retention time using IntantPC-labeled glycan standards whenever available. Agilent OpenLab ChemStation™ software produced a sequence report that included glycan identity, summary (retention time and peak area), and chromatograms.

#### Microfluidic modulation spectroscopy (MMS)

The secondary structure (i.e., folding of the protein backbone) of LF was measured using microfluidic modulation spectroscopy (MMS) with an AQS^3^pro instrument from RedShiftBio (Boxborough, MA, USA). Dialysed samples in 50 mM Tris (pH 7.5) were loaded in triplicate into a 24-well plate in sample/buffer pairs and were run on the AQS^3^pro MMS system that was equipped with sweep scan functionality and a flow cell with a 22.3 µm path length. An average of eight spectra were collected with a 5 psi backing pressure scan across the amide I band between 1588 and 1712 cm^-1^. All samples were analyzed using the AQS^3^pro delta Data Analysis software (version 2.5.98.4131). Lysozyme was used as the model protein. The samples were fit to the region between 1720 and 1680 cm^-1^. The nominal displacement factor was set to 0.6, the second derivative smoothing window was set to 19 cm^-1^, and the baselining setting (subtraction) was “Rubberband.” A higher-order (secondary) structure was calculated using “Globular protein” pre-settings for the Gaussian curve deconvolution.

#### Iron saturation

The iron content of the rhLF and hmLF protein samples was measured according to the AOAC-2015.01 method. Iron saturation was calculated with the following formula:

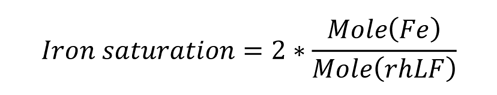

*Mole (Fe)* and *Mole (LF)* were the moles of iron atoms and LF, respectively. They were calculated using the measured iron content and LF protein content with a MW of 56 for iron and 80,000 for LF.

#### Small-angle X-ray scattering (SAXS)

Lactoferrin samples were prepared by dialysis in 50 mM Tris with 150 mM NaCl (pH 7.5) using 10,000 Da MW cut-off Slide-A-Lyzers^TM^ (ThermoFisher). Lactoferrin protein samples were loaded onto a quartz capillary flow cell at 4 °C and precisely aligned with the X-ray beam under vacuum conditions (< 1×10⁻³ torr). Small-angle X-ray scattering (SAXS) data was gathered in triplicate at concentrations of 1.0 and 4.0 mg/mL. A Rigaku BioSAXS-2000 nano SAXS Kratky camera system (The Woodlands, TX, USA) was used to conduct the experiment. A Rigaku MM007 rotating anode generated X-rays with a wavelength of 1.5418 Å. The system was equipped with OptiSAXS confocal max-flux optics with a HyPix-3000 Hybrid Photon Counting detector. The useful q-space range, where *q* represented the scattering vector (4πsinθ/λ, with 2θ being the scattering angle), typically spanned from *q_min_* = 0.008 Å⁻¹ to *q_max_* = 0.6 Å⁻¹. The X-ray beam had an energy of 8.04 keV, with a Kratky block attenuation of 22% and a beam diameter of approximately 100 μm. A total of six 10-min images were collected and averaged. Obtained SAXS data underwent buffer subtraction, resulting in the final dataset we used for subsequent analysis. Data processing, including image integration, normalization, and background buffer data subtraction, was executed using Rigaku SAXSLAB software.

#### Analytical Ultracentrifugation (AUC)

AUC analysis was carried out with an Optima multiwavelength instrument (Beckman Coulter, Indianapolis, IN) equipped with absorbance and interference optics. Samples were equilibrated to 20 °C. The instrument was operated under a full vacuum and the An-50 titanium rotor was spun at 30,000 RPM for 16.5 hr; radial scans of the cells were recorded every two min at 280 nm. AUC data was analyzed with UltraScan III [45, 46]. Reference scans were automatically selected to convert the data from raw radial intensity to pseudo-absorbance. The air-liquid meniscus was then manually selected for each sector. The data range was also manually selected (usually between 6.0 cm and 7.1 cm). The first five to 10 scans were excluded from analysis, as were scans at the end with little to no sedimentation signal. Data was fitted with an S-value range between 1 and 10, with a resolution of 100, a frictional ratio range of 1 and 4, and a resolution of 64. Time invariant noise was fitted during the first two-dimensional spectral analysis. When residuals were <0.003, data was re-fitted for time and radially invariant noise, and the meniscus was adjusted. When the correct meniscus was identified, an additional time and radial invariant noise fit was performed using an iterative fitting method with a maximum of 10 iterations. The resulting pseudo-three-dimensional plots were evaluated for final S-values, frictional ratios, and MW calculations.

#### Differential scanning calorimetry (DSC)

Differential scanning calorimetry (DSC) was carried out using MicroCal VP-capillary DSC (Malvern Panalytical, Westborough, MA, USA). Data was collected over a temperature range of 25 °C to 105 °C with a scan speed of 90 °C/hr. A buffer reference experiment was conducted for baseline subtraction. The cell was loaded with 400 µL of sample at a concentration of 2.0-5.0 mg/mL in the buffer 50 mM tris at pH 8.0. The running software was VPViewer2000. Raw data was converted into molar heat capacity using Microcal, LLC Cap DSC Version Origin70-L3.

## RESULTS and DISCUSSION

The objective of this study was to conduct a thorough characterization and analysis of Helaina rhLF protein produced at an industrial scale using a *K. phaffii* fermentation process. Various analytical techniques were employed to confirm identity, assess purity and impurities, measure size, and investigate the structure of Helaina rhLF protein, including glycosylation (Table 1). Additional comparisons to hmLF and bLF samples were made.

**Table 1.** Analytical techniques exploited to analyze and characterize Helaina rhLF as well as native hmLF.

| Analytical technique | Attributes of protein |
| --- | --- |
| SDS-PAGE/Western blotting | Molecular weight / Identity |
| Reversed-phase high performance liquid chromatography (RP-HPLC) | Protein content and purity |
| HPLC-fluorescence detection | Released glycan profile |
| ICP-MS | Iron content and<br>Iron saturation (in conjunction with HPLC data) |
| Mass spectrometry (including LC-MS) | Protein sequence<br>Posttranslational modifications<br>Glycosylation<br>Host cell proteins |
| Microfluidic modulation spectroscopy (MMS) | Secondary structure |
| Analytical Ultracentrifugation (AUC) | Tertiary structure and aggregation/degradation |
| Small Angle X-Ray Scattering Spectroscopy (SAXS) | Tertiary structure and aggregation/degradation |
| Differential scanning calorimetric (DSC) | Structure and thermostability |

### Protein identity

Helaina rhLF was designed according to the protein ID of hLF, specifically TRFL_HUMAN (Uniprot P02788) of the UniProt database. Like mature hmLF, Helaina rhLF did not have the N-terminal signal peptide (i.e. amino acid sequence 1-19), as shown in the theoretical sequence (Fig. 2A). Helaina rhLF had a total of 691 amino acid residues with the N-terminal starting with the residue glycine (G)-19 and C-terminal ending with lysine (K)-710.

**Fig. 2(A).**
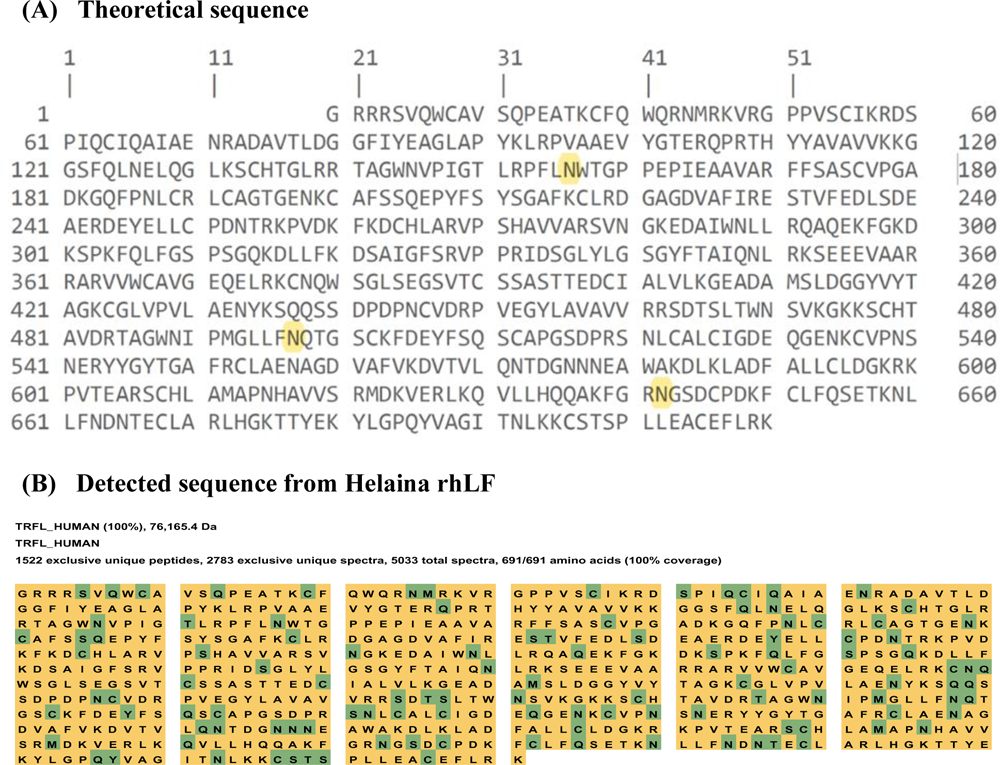
Protein sequence of native hmLF (Uniprot P02788) without the N-terminal signal peptide (amino acid sequence 1-19). The highlighted N indicates the three known N-glycosylation sites, namely, Asn-156, -497 and -642; (B). Detected sequence of native hmLF from Helaina rhLF as highlighted by the yellow and green color. The green color indicates posttranslational modifications.

The nanoflow LC-MS/MS-based bottom-up proteomics approach was used to determine the amino acid sequence of Helaina rhLF. In this study, we used trypsin, chymotrypsin, and elastase to maximize the sequence coverage [47, 48]. Trypsin specifically cleaved at the C-terminal side of lysine and arginine residues, while chymotrypsin and elastase cleaved non-specifically at the hydrophobic amino acid residues alanine, isoleucine, leucine, and valine. The combination of trypsin, chymotrypsin, and elastase produced complementary peptide sequences that covered the entire sequence of Helaina rhLF. Proteomics analysis led to the identification of Helaina rhLF as TRFL_HUMAN with 100% sequence coverage (Fig. 2B). Protein identification was based on detecting more than 1200 unique peptides generated by proteolytic digestion, in conjunction with the stringent criteria for peptide sequence identification as described in the Materials and Methods section. The data confirmed that the amino acid sequence of Helaina rhLF was identical to that of the hmLF. Additionally, SDS-PAGE and western blotting data showed that Helaina rhLF had a similar MW to that of hmLF and was positively detected with anti-hLF antibody (Fig. 3), further confirming that the size and identity of Helaina rhLF were identical to hmLF.

**Fig. 3.**
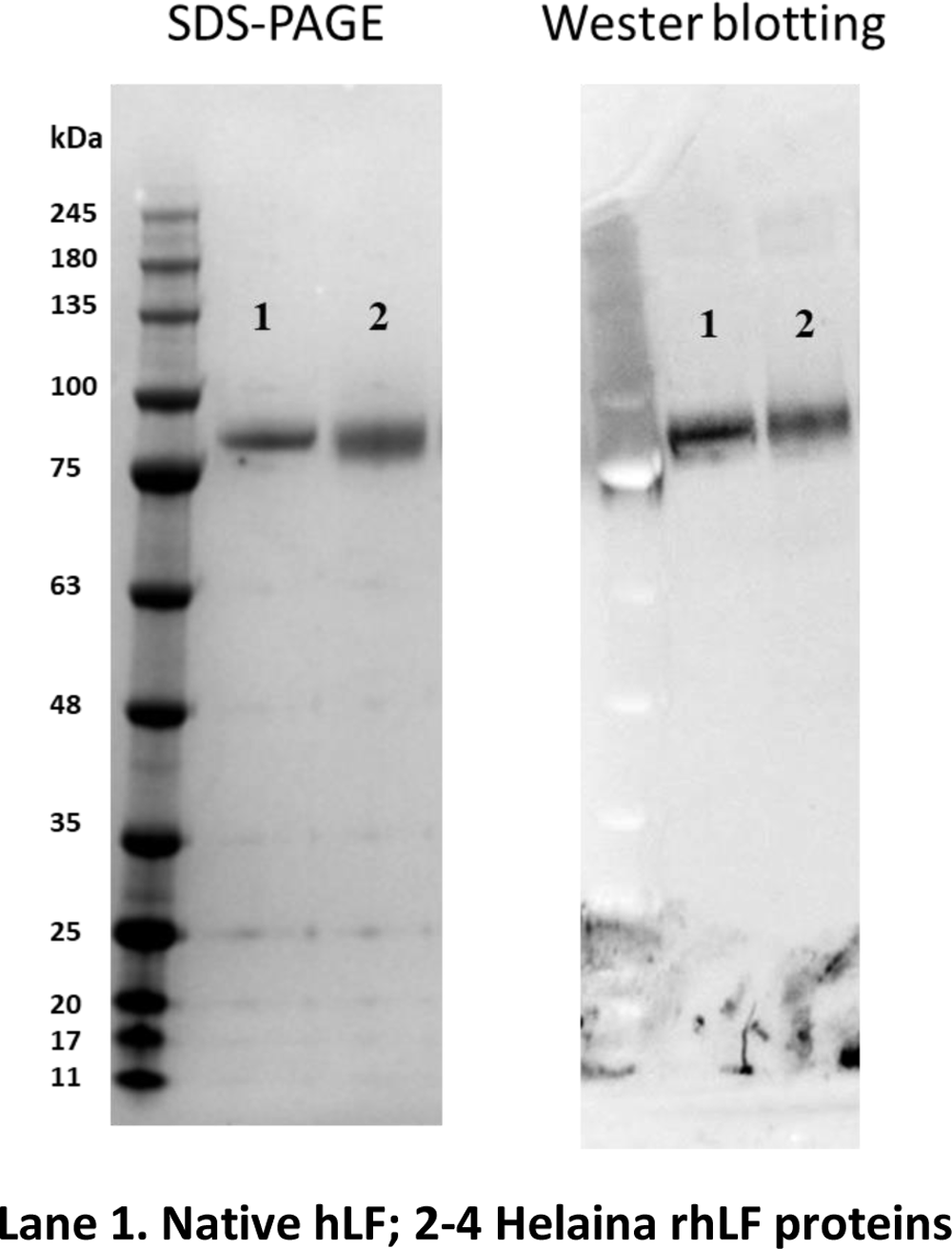
Representative images of SDS-PAGE and western blotting of native hmLF (Lane-1) and Helaina rhLF (Lane-2). Other conditions are described in Materials and Methods.

### Protein purity

Helaina rhLF produced from the *K. phaffii* recombinant system underwent multiple isolation steps by microfiltration/diafiltration followed by cation exchange chromatography as the final purification step. Isolated rhLF protein purity was assessed by multiple analytical techniques including SDS-PAGE, RP-HPLC with UV detection, and nanoflow LC-MS/MS-based proteomics approach.

Fig. 3 shows a representative SDS-PAGE image of 300 ng of Helaina rhLF and hmLF. Colorimetric staining with a limit of detection 3 ng, detected no visible protein bands, suggesting that protein contaminants, if any, in Helaina rhLF, were less than 1%. RP-HPLC analysis provided an orthogonal measurement of Helaina rhLF quantification and purity. As shown in Fig. 4, Helaina rhLF was detected as the predominant peak along with a couple of minor peaks at both wavelengths of 280 nm (more specific to protein) and 214 nm (less specific to protein). The average purity of three batches of rhLF was 98% with a small relative standard deviation of 0.4%, based on peak area percentage calculations obtained at 214 and 280 nm. Further LC-MS/MS-based proteomics analysis of the Helaina rhLF revealed that identified proteins were predominantly human LF with an average relative abundance >99%. No residual host cell proteins (HCP) had a relative abundance greater than 0.1%, with the top three identified HCP being phosphoglycerate kinase (UniProt i.d. C4QY07, 0.04%), endoplasmic reticulum chaperone BiP (UniProt i.d. C4QZS3, 0.04%), and SCP domain-containing protein (UniProt i.d. C4R3H3, 0.03%) (see supplementary Table S1). It should be noted that MannosidaseTR is not a residual protein from the yeast expression system but rather a protein introduced in the engineering process to help minimize hyper-mannosylation of yeast.

**Fig. 4.**
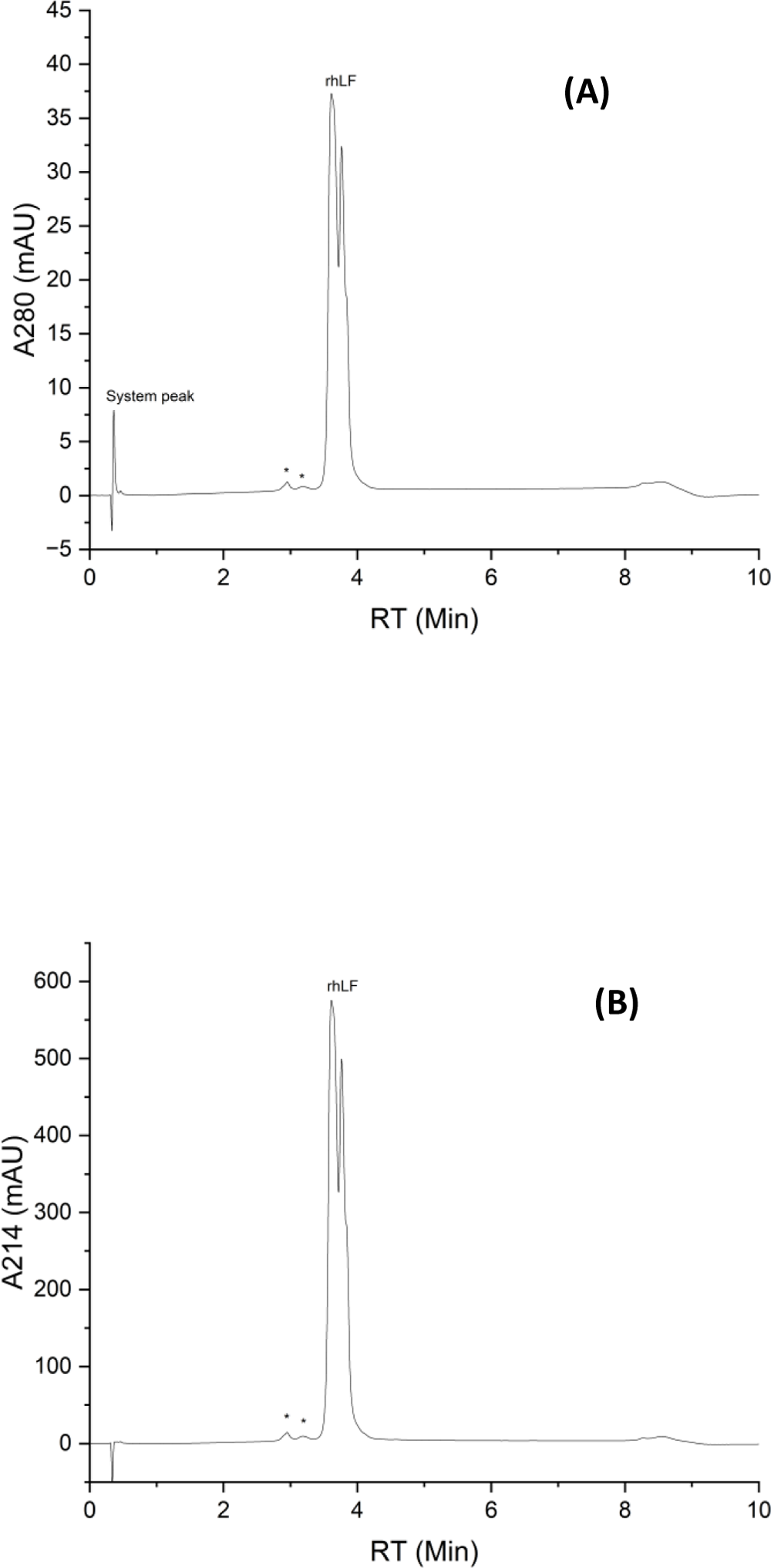
Representative chromatograms of RP-HPLC analysis of Helaina rhLF with UV detection at 280 nm (A) and 214 nm (B). * Indicates impurity peaks observed from the rhLF samples. The chromatograms were generated from 1.0 µL injection of 1.0 mg/mL Helaina rhLF. Other conditions are described in Materials and Methods.

### Glycosylation

Two complementary analytical techniques; site-specific glycosylation analysis and released glycan profiling, were employed to gain a full understanding of the glycosylation of Helaina rhLF protein.

Site-specific glycosylation properties of LF were revealed by the LC-MS/MS-based proteomic analysis. N-glycans were detected at all three predicted asparagine (Asn) sites as described for the hmLF, Asn-156, Asn-497 and Asn-642 [11, 49]. In Helaina rhLF, all three asparagine sites share the same set of N-glycans-oligomannose N-glycans containing 5 to 14 mannose residues (M5-M14) (Fig. 5). No complex or hybrid types of N-glycans were identified. The occupancy rates for the three sites, however, were different and none of the three sites were completely occupied by glycans. The occupancy rate was 75% for the site Asn-156; 80% for Asn-497; and 18% for Asn-642, respectively (Fig. 5). These observations were similar to the results reported by Zlatina et. al., where there were approximately 90% occupancy rates for both Asn-156 and Asn-497, but only 9% for Asn-624, respectively [12].

**Fig. 5.**
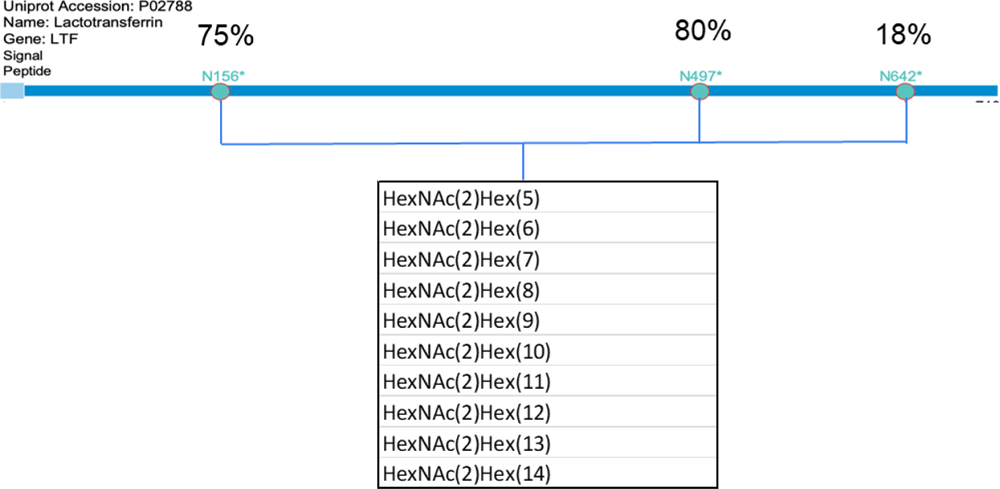
The identified N-glycans as well as the glycan occupancy rates on the three glycosylation sites (i.e. Asn-156, -497 and -642) of Helaina rhLF.

Fig. 6A shows a representative chromatogram of the released N-glycan profile of Helaina rhLF protein and Table 2 lists the relative abundance of the identified glycans, estimated based on their peak areas. Same as the results from the site-specific analysis, the identified N-glycans were primarily oligomannoses M5-M14. The smaller oligomannose M5-M9 accounted for about 59±2% of total identified N-glycans while the relatively larger oligomannose M10-M14 only took 41±2% (Table 2). Interestingly the mass spectrometric data (not shown) also suggested there were about 23% phosphorylated M5-M14. The phosphomannose glycans present on Helaina rhLF are the same oligomannose structures (M5-M14) with either 1 or 2 phosphate groups. Phosphomannose structures have been observed in numerous recombinant glycoproteins produced in *K. phaffii*, in abundances ranging from 20% to 30%, although their presence depends on the protein being expressed [50–53]. This range of abundance coincides with that measured in Helaina rhLF. The results of the N-glycan profiles of Helaina rhLF were highly consistent across three different batches with an average relative standard deviation (RSD) of the relative abundance of 9% and a range of 2%-12% (Table 2 and Fig. S1).

**Fig. 6.**
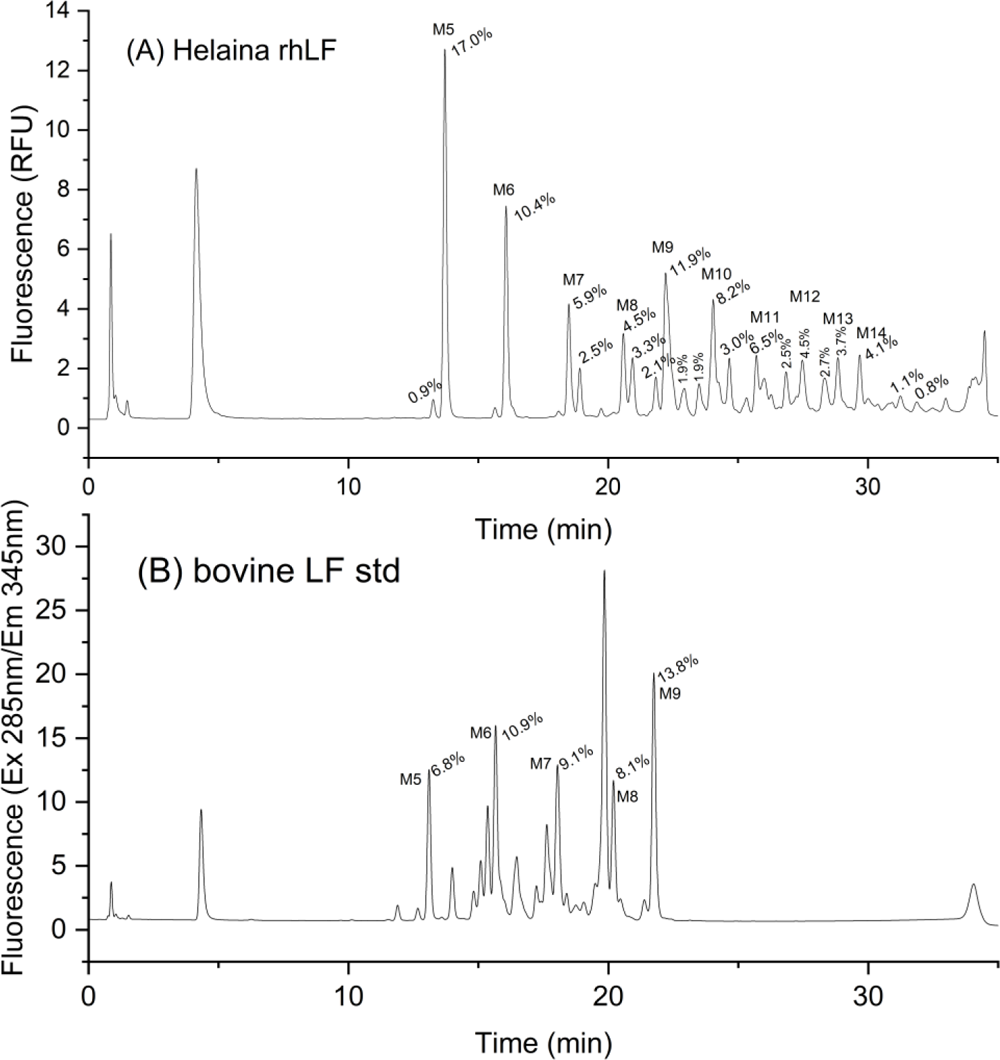
Comparison of the released N-glycan profiles of Helaina rhLF (A) and bLF (B). Conditions for data acquisition and processing were detailed in the Material and Methods.

**Table 2.**
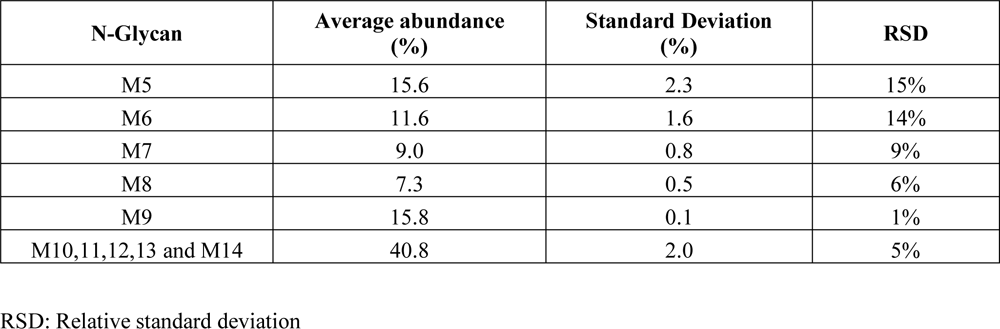
The average relative abundance (%) and reproducibility of the N-glycans identified from three batches of Helaina rhLF.

| N-Glycan | Average abundance (%) | Standard Deviation (%) | RSD |
| --- | --- | --- | --- |
| M5 | 15.6 | 2.3 | 15% |
| M6 | 11.6 | 1.6 | 14% |
| M7 | 9.0 | 0.8 | 9% |
| M8 | 7.3 | 0.5 | 6% |
| M9 | 15.8 | 0.1 | 1% |
| M10,11,12,13 and M14 | 40.8 | 2.0 | 5% |

RSD: Relative standard deviation

Fig. 6B shows that the smaller oligomannoses M5-M9 were detected in bovine milk LF at high abundance along with complex and hybid glycans. It should also be pointed out that M4-M9 have been reported in hmLF by different labs [15, 54] In our study, we consistently detected M5-M6 in hmLF from different sources including those supplied by MilliporeSigma and Abcam as well as purified from breast milk by Helaina (Fig. 7 and S2).

**Fig. 7.**
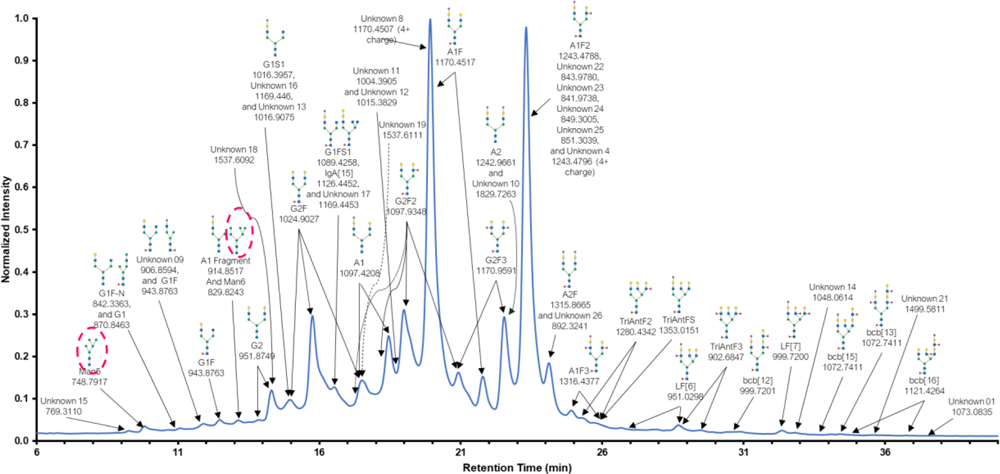
Representative LC-MS chromatogram of the released N-glycan profile of native hmLF. The detected oligomannose M5 and M6 are indicated with red circles. Each peak is marked with the observed m/z in the positive mode and assigned with the best matching glycan structures. Other conditions are described in Materials and Methods.

### Protein structures

#### Secondary structure by MMS

The secondary structure of Helaina rhLF was assessed using MMS and then compared to secondary structure of hmLF and bLF. MMS is a novel infrared (IR) technology that has been gaining increasing interest for protein structural analysis and similarity comparison, due to its simplicity, sensitivity, and automation capabilities [55, 56]. MMS monitors the vibrational response of the amide I band at 1600 - 1700 cm^-1^, representing the stretching of the carbonyl group of the amide bonds. The amide I vibrational signals are sensitive to the H bonding environment and the folding of the protein backbone (the protein secondary structure).

Fig. 8A shows representative second derivative MMS spectra of Helaina rhLF, hmLF, and bLF. They all comprise the same types of secondary structure features, namely α-helix, β-sheet, and turn, as evident by the observations of the distinct absorbance peaks at 1656, 1634, and 1686 cm^-1^ [57, 58]. Helaina rhLF’s MMS spectrum overlaps exactly with that of hmLF but not that of bLF. Fig. 8B further compares the compositions of the secondary structures of Helaina rhLF (averaged from three batches) with hmLF (averaged from three hmLF from three sources). Helaina rhLF comprises 38%±1% α-helices, 28%±1% β-sheets, and 23%±1% turns versus hmLF’s 40%±1% α-helices, 29%±1% β-sheets, and 23%±1% turns. This data suggests Helaina rhLF secondary structure was nearly identical to that of hmLF. The MMS results of Helaina rhLF and hmLF closely match previously published data for native hmLF as measured. The MMS results of Helaina rhLF and hmLF closely match previously published data for native hLF as measured by circular dichroism (CD); where 37.4% α-helices, 28.2% β-sheets, and 15.8% turns were reported [59]. Additionally, the CD measurements of rhLF expressed in transgenic mice showed similar results with 40% α-helices, 20% β-sheets, and 18% turns [60].

**Fig. 8.**
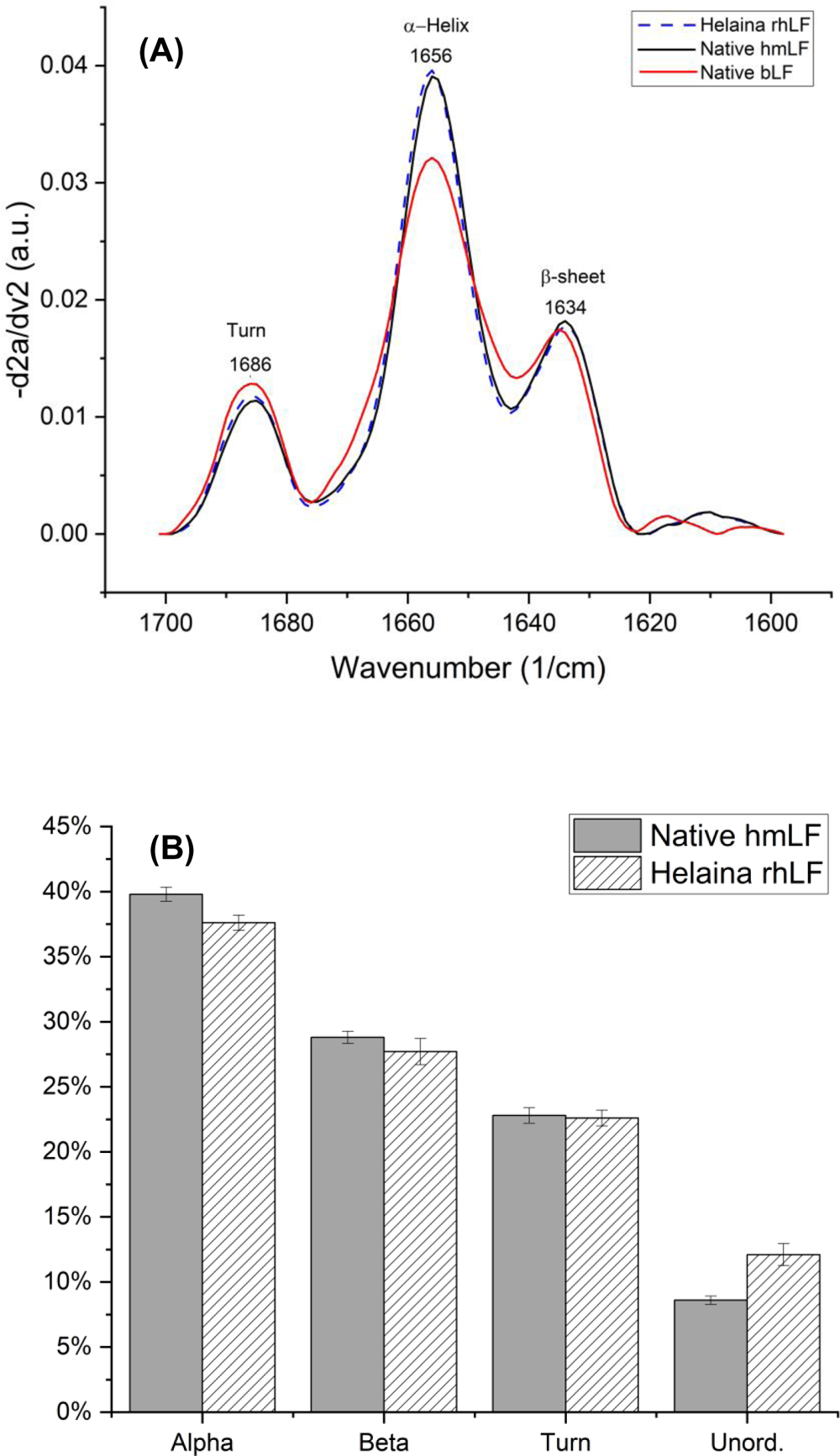
Representative second derivative MMS spectra of Helaina rhLF, native hmLF and native bLF (A); and comparison of the compositions of the secondary structures of Helaina rhLF with native hmLF (B). The composition of the secondary structures of Helaina rhLF was the average from three production batches, while that of native hmLF was the average from three different sources of hmLF as described in the Material section.

Similarity scores [61] are useful to quantify the degree of structural similarity between proteins. We computed similarity scores by normalizing the overlap of the second derivative MMS spectra of a LF protein (hmLF or Helaina rhLF) with the hmLF from MilliporeSigma, according to the approach proposed by Kendrick et al [61]. The mean similarity score was 98.2%±1.6% (range 96%-100%) for hmLF from three different sources, and 96.2%%±1.4% (range 95%-98%) for three Helaina rhLF batches, respectively. The hmLF from MilliporeSigma had a similarity score of 100% since it was used as the reference for the similarity score computation for all the LF. Nevertheless, the range of the similarities score of hmLF and Helaina rhLF overlapped and the difference in the means was only 2.0%. This provides further support that Helaina rhLF’s MMS secondary structures were nearly identical to those of hmLF. The nearly identical secondary structures between Helaina rhLF and hmLF can be attributed to their identical amino acid sequences as determined by the proteomics analysis, since the secondary structure of a protein is predominantly determined by its primary structure.

#### Iron saturation

As an iron-binding protein, each LF molecule has two lobes, each capable of binding one iron atom. Iron saturation reflects the percentage of two lobes filled with iron atoms. The mean (standard deviation) iron saturation rate of three batches of Helaina rhLF was 54% (±4%) and ranged between 50% and 60%. These results indicate that about one-half to two-thirds of the lobes of Helaina rhLF contained iron. By way of comparison, hmLF is typically the apo-form, meaning its saturation was under 20% [62].

#### Tertiary structure by SAXS and AUC

We used SAXS to determine and compare the tertiary structure of Helaina rhLF to that of hmLF and bLF. The SAXS analysis reveals information about the size, shape, and conformational flexibility of proteins in solution [63–66]. We then utilized the reported crystallographic structures of hmLF to validate the accuracy of the molecular envelopes reconstructed from SAXS. Additionally, we employed AUC to further validate the structural information obtained by SAXS.

SAXS raw data (Fig. S3) for Helaina rhLF, hmLF, and bLF was acquired at 1.0 mg/mL (A) and 4.0 mg/mL (B) concentrations. The SAXS data sets were collected for 60 min, with six 10-min images for each concentration. The overlays of these images were averaged and showed there was no X-ray radiation damage. Replicate datasets were also averaged. A comparison of the SAXS data between low and high concentrations showed near-identical signal profiles and no concentration-dependent multimerization. As expected, compared to the lower concentration, the higher concentration produced a greater signal to noise SAXS data.

Raw data (Fig. S3) was reconstructed into Guinier plots (Fig. S4), Pair distance distribution function P(r) plots (Fig. S5), and Kratky plots (Fig. S6), molecular envelopes (Fig. 9) to gain the understanding of the characteristics of the size, shape, and conformation of the LF proteins. The results are summarized in Table 3.

**Fig. 9.**
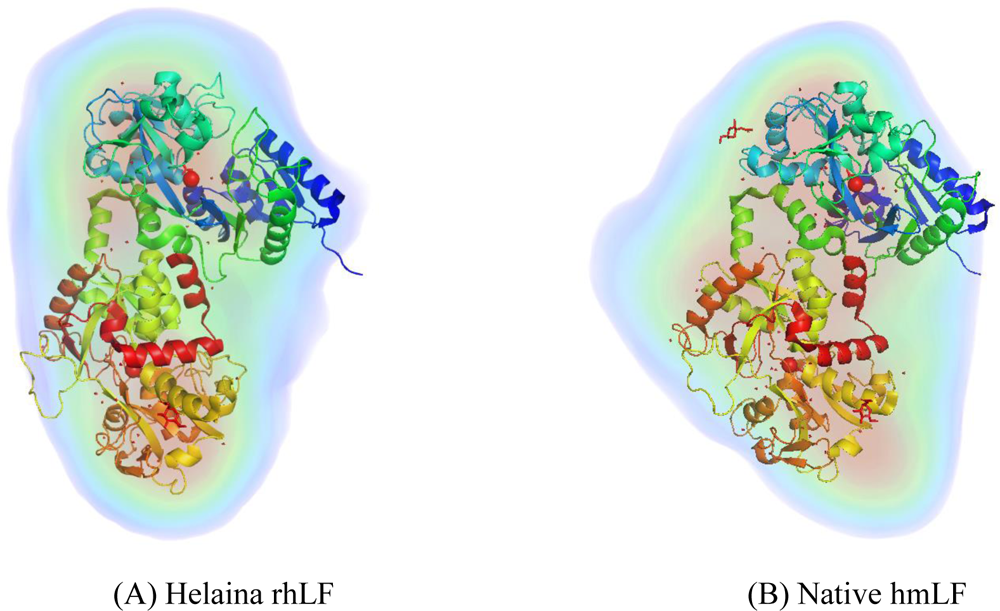
Representative solvent envelopes of Helaina rhLF and native hmLF derived from the SAXS data and their fits with the Fe^+3^ bound human lactoferrin monomer (2bjj) model. Conditions for data acquisition and processing were detailed in the Material and Methods.

**Table 3.** Characteristics of LF proteins revealed by AUC and SAXS analyses.

| LF protein | f/f0 (AUC) | Aggregate d | Guinier $R_g$ (Å) | P(r) $R_g$ (Å) | $D_{max}$ (Å) | Porod Volume (Å<sup>3</sup>) | SASREF Chi-sq fit to holo-hmLF | SASREF Chi-sq fit to apo-hmLF |
| --- | --- | --- | --- | --- | --- | --- | --- | --- |
| *Helaina rhLF* |  |  |  |  |  |  |  |  |
| Mean | 1.32 | No | 41.3 | 41.5 | 161.0 | 1.3E+05 | 3.2 | 3.4 |
| STDEV | 0.04 |  | 13.37 | 13.59 | 83.71 | 3.6E+04 | 1.79 | 2.33 |
| RSD | 2.7% |  | 32.4% | 32.7% | 52.0% | 26.7% | 56.4% | 67.6% |
| *Native hmLF* |  |  |  |  |  |  |  |  |
| hmLF-S | 1.31 | No | 35.4 | 35.5 | 122.5 | 1.2E+05 | 3.1 | 2.9 |
| hmLF-A | 1.29 | Yes | 56.6 | 57.1 | 257 | 1.8E+05 | 5 | 6 |
| hmLF-H | 1.36 | No | 31.9 | 32 | 103.4 | 1.1E+05 | 1.42 | 1.44 |
| Mean | 1.32 |  | 41.3 | 41.5 | 161.0 | 1.3E+05 | 3.2 | 3.4 |
| STDEV | 0.04 |  | 13.37 | 13.59 | 83.71 | 3.6E+04 | 1.79 | 2.33 |
| RSD | 2.7% |  | 32.4% | 32.7% | 52.0% | 26.7% | 56.4% | 67.6% |

The plots of Guinier fit (Fig. S4) and pair distance distribution function P(r) (Fig. S5) display the overall size and multimerization/aggregation of the LF proteins in solution (50 mM tris with 150 mM sodium chloride, pH 7.5 for our analysis). The Guinier fit determined the radii of gyration (*R_g_*) of LF proteins (Table 3). No aggregation was observed for all tested LF proteins, except for the hmLF-A (hmLF from Abcam) and bLF-T (bLF from The Lactoferrin Company) proteins. As can be seen, Helaina rhLF and the hmLF proteins, except hmLF-A, had *R_g_* had values between 32 and 35 Å, which was in agreement with the size of a protein monomer from the X-ray crystal structure model of hmLF (PDB codes 2BJJ and 1LFH). Fig. S4 shows a “smiley Guinier” fit with a radius of gyration, *R_g_* of 42.8 (± 0.3) Å for the hmLF-A and 110.7 (± 0.6) Å for bLF-T, indicating protein aggregation. Fig. 10 is a comparison of the *R_g_* of Helaina rhLF and native hmLF proteins, showing similar average *R_g_* with an overlapping distribution. The P(r) plot (Fig. S5) is a histogram of the distances between pairs of points within the LF proteins in solution. Rmax, where the P(r) curve intersects the X-axis, is the maximum diameter of the proteins. All Helaina rhLF and hmLF and bLF, except hmLF-A and bLF-T, appeared to be globular and peaked around 35 Å with a diameter of 128 Å in the range of the *R_g_* values from the Guinier analysis. The hmLF-A and bLF-T proteins had broad humps following the main peak and corresponding to the large multimers with a maximum diameter over 250 Å, respectively. These results indicate that both Helaina rhLF and hmLF were present as monomers in aqueous solution and with similar overall size.

**Fig. 10.**
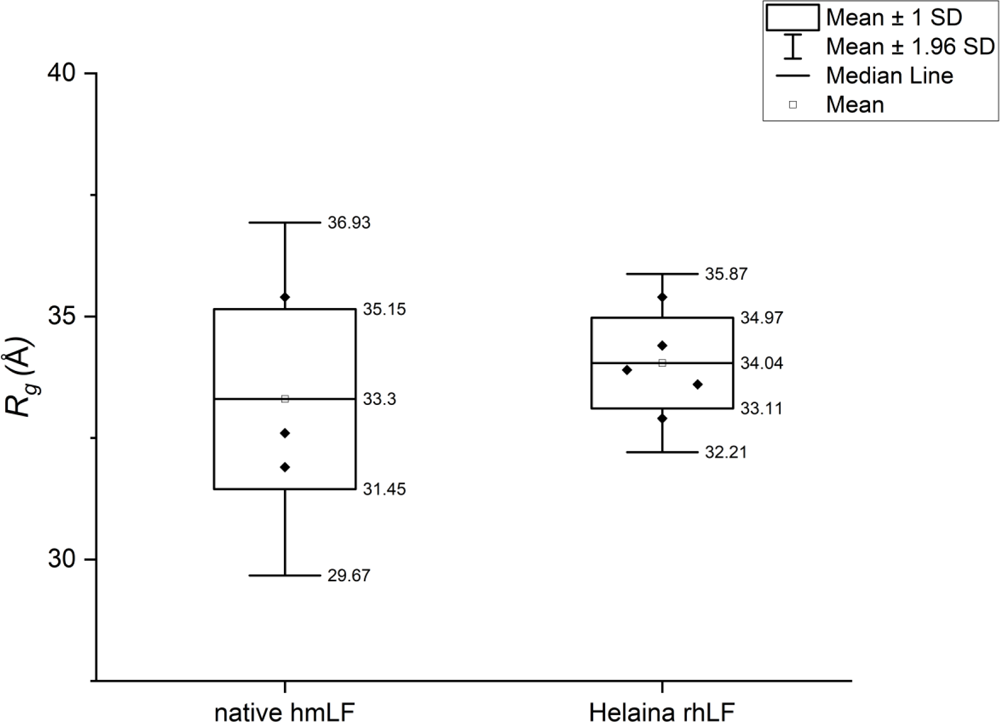
Comparison of the radii of gyration (*Rg*) of the Helaina rhLF proteins with native hmLF proteins. Conditions for data acquisition and processing were detailed in the Material and Methods.

Kratky plots derived from SAXS data provided a further qualitative assessment of the flexibility and/or degree of unfolding in the tested LF protein samples (Fig. S6). Proteins that are unfolded or highly flexible typically exhibit a plateau in the Kratky plot at high scattering vectors (s), while compact, globular proteins display a bell-shaped Gaussian peak. The Kratky plots we generated indicate that all tested human and bovine LF were well-folded globular proteins with no discernible flexibility or disorder.

Fig. 9 illustrates the solvent envelopes of Helaina rhLF and hmLF derived from SAXS data. These envelopes closely matched the reported X-ray crystal structures of both Fe^+3^ bound (holo) and iron-free (apo) hmLF monomers (2BJJ and 1LFH in the RCSB PDB database, respectively). The fits were rigorously validated through refinement using normal mode analysis in Sreflex, leveraging SAXS data. The Pymol-generated representation incorporated a transparent surface overlying the electron density from solution scattering (DENSS), the latter being an advanced algorithm for direct computation of *ab initio* electron density maps using solution scattering data.

The SAXS refinement, employing the SASREF method, successfully refined the N and C-lobes of LF individually, resulting in improved Chi-square values and better fits into the DENSS envelopes. As indicated by the SASREF Chi-square values (Table 3), all Helaina rhLF and hmLF had a better fit to the holo monomer than to the apo monomer, with the exception of hmLF-S (from MilliporeSigma). An explanation for this discrepancy may be that Helaina rhLF was not in an apo form but rather 50%-60% iron-saturated, while hmLF-S was an essentially apo form with approximately 8% iron saturation.

The AUC results (Fig. 11 and S7) also demonstrated the consistency of Helaina’s rhLF industrial-scale production process: Fig. S7A-C, showing the three batches of rhLF in comparison to hmLF-S (from MilliporeSigma, Fig. S7D), and hmLF-H (purified in house, Fig. S7F). The hmLF-A (from Abcam) showed multiple species, suggesting more heterogeneity than the recombinant batches or the other native samples (Fig. S7E). Each recombinant batch had highly similar frictional ratios of 1.25 – 1.30, suggesting a globular protein. This correlated with the crystal structure of hmLF and with the SAXS results. Furthermore, MW estimates from each batch were 76 – 80 kDa, and appear to be in agreement with the expected MW of native hLF (∼ 80 kDa) and with the MW estimates obtained for hmLF-S (from MilliporeSigma) and hmLF-H (purified in house) (Fig. S7D and F).

**Fig. 11.**
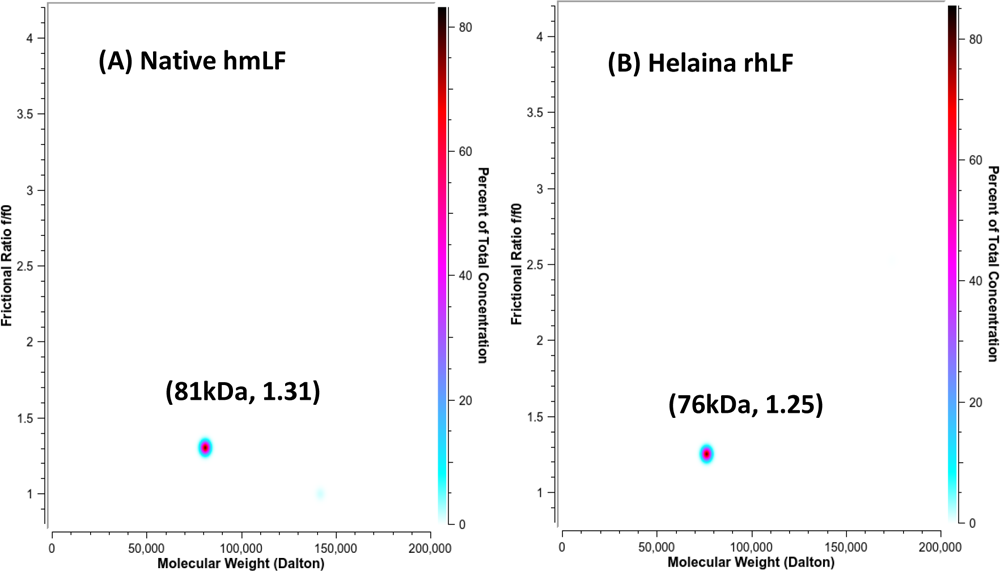
AUC measurements of native hmLF (A) and Helain rhLF (B). The data presented a pseudo-three-dimensional distribution of the observed protein species, with the calculated molecular weight on the x-axis, the calculated frictional ratio on the y-axis, and the % concentration in the Z-plane, with the heat map on the right axis. Conditions for data acquisition and processing were detailed in the Material and Methods. Conditions for data acquisition and processing were detailed in the Material and Methods.

In summary, both Helaina rhLF and hmLF were confirmed as monomeric, well-folded, globular proteins. The slight variations in the shapes of their molecular envelopes can be attributed, in part, to the flexibility in the orientation of the N-lobe and C-lobe domains under various conditions, particularly in response to changes in iron saturation.

#### Thermostability by DSC

Protein unfolding under heat is primarily determined by the protein’s structure, and the more tightly a protein is folded, the higher temperature that is required to unfold (i.e., denature). We employed DSC to compare the thermal stability of Helaina rhLF and hmLF. Fig. 12A shows the hmLF as isolated, which comprises predominantly the apo form with a low (∼8%) iron saturation, underwent denaturation (unfolding) at maximum peak temperature of approximately 67 °C. The holo-, iron-saturated (∼99%) form of hmLF exhibited drastically higher thermal stability than the apo form with a much-elevated denaturation temperature of approximately 91 °C. Helaina rhLF had two denaturation temperatures, 68.7±0.9 °C and 91.9±0.5 °C (Table 4 and Fig 12), due to Helaina rhLF’s partial iron saturation (50-60%). In other words, Helaina rhLF was a mix of both apo-and holo-rhLF. Helaina rhLF having identical denaturation temperatures to that of apo and holo hmLF provided further evidence that they have highly similar secondary and tertiary structures.

**Fig. 12.**
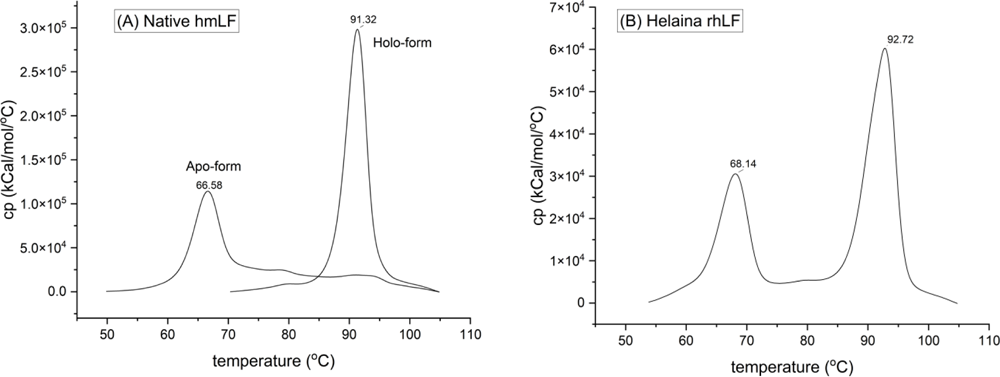
Comparison of differential scanning calorimetry of native hmLF (apo-and holo-form) (A) and Helaina rhLF (B). Conditions for data acquisition and processing were detailed in the Material and Methods.

**Table 4.** Maximum peak temperature (T_max_) of Helaina rhLF and native hLF.

| | $T_{\max 1}$ (°C) | $T_{\max 2}$ (°C) |
| --- | --- | --- |
| Helaina rhLF | 68.7±0.9 | 91.9±0.5 |
| Native hmLF-Apo | 66.4±0.5 |  |
| Native hmLF-Holo |  | 91.1±0.5 |

## CONCLUSION

In summary, our study presents a comprehensive analysis of industrial scale batches of Helaina rhLF (Effera™) expressed in the *K. phaffii* system. We showed that Helaina rhLF has a protein purity of 98% or higher, confirmed through RP-HPLC, SDS-PAGE, and LC-MS/MS-based proteomics. LC-MS/MS analysis verified the 100% identical amino acid sequence to hLF, with N-linked glycans detected at the same sites found on hmLF. The analysis has also identified that the N-glycans in Helaina rhLF are oligomannose structures (M5-M9), with M5 and M6 detected in hmLF samples, although complex N-glycans are the predominant ascribed glycoforms. MMS revealed that the secondary structure of Helaina rhLF closely mirror those of hmLF, with an average 96% similarity that can be attributed to identical amino acid sequences. Additionally, data generated from the SAXS, AUC, and thermostability analyses confirmed that Helaina rhLF structure closely resembles hmLF, exhibiting well-folded globular structures and undergoing denaturation at temperatures consistent with apo-and holo-hmLF. Overall, these findings demonstrate that Helaina rhLF is of high purity and has a structure akin to hmLF.

## Acknowledgements

The authors thank Julia Fecko of the X-ray Crystallography core at the Penn State University, Huck Institutes of the Life Sciences, for her invaluable assistance in the DSC analyses. Additionally, support for small-angle X-ray scattering was provided by SIG S10 of the National Institutes of Health under award number # S10-OD028589 to N.H.Y. The Optima AUC utilized in this research was made possible by NIH grant S10 OD032215-01to N.H.Y. The authors would also like to thank Drs. Sheng Zhang and Qin Fu at the Proteomics and Metabolomics Facility of Cornell University for providing the mass spectrometry data and NIH SIG grant 1S10 OD017992-01 support for the Orbitrap Fusion mass spectrometer. The authors thank Dr. Valerie Collins and Rich DiVirgilio at RedShiftBio for help in the MMS analyses and Julie Wushensky, Dr. Greg Kilby, and Chuck Cloyd at MOBILion Systems, Inc for assistance in the glycan analyses. The authors want to specially thank Carolyn Alish for her assistance in writing, editing, and proofreading the manuscript.

