## Supplemental data for "Characterization of Recombinant Human Lactoferrin Expressed in *Komagataella Phaffii*"

#### Supplementary Data

##### Table S1. The relative abundance (%) of the identified proteins from Helaina rhLF

| Identified Proteins | Accession Number | Helaina rhLF-1 | Helaina rhLF-2 | Helaina rhLF-3 | Average |
|---|---|---|---|---|---|
| LACTOFERRIN | LACTOFERRIN | 99.76877025 | 99.66679381 | 99.92559697 | 99.78705368 |
| Phosphoglyceratekinase | C4QY07 | 0.071008065 | 0.055368055 | 0.010088388 | 0.045488169 |
| Endoplasmic reticulum chaperone BiP | C4QZS3 | 0.03425931 | 0.060051426 | 0.034198497 | 0.042836411 |
| MannosidaseTR | MannosidaseTR | 0.020111374 | 0.074992832 | 0.003290942 | 0.032798383 |
| SCP domain-containing protein | C4R3H3 | 0.026875124 | 0.043759254 | 0.014924343 | 0.028519573 |
| Catalase | C4R2S1 | 0.008502079 | 0.024912314 | 0.008422246 | 0.013945546 |
| 5-methyltetrahydrop teroyltriglutamate homocysteine S-methyltransferase | C4QZU2 | 0.009931174 | 0.012304305 | 0 | 0.011117739 |
| Superoxidedismutase [Cu-Zn] | C4R8X7 | 0.005425298 | 0.023314325 | 0.001151436 | 0.009963687 |
| Ribonuclease T2 | C4QZY8 | 0.012172773 | 0.008977466 | 0.003208633 | 0.008119624 |
| Cytochromec isoform 1 | C4R6L9 | 0.015564249 | 0.000428016 | 0 | 0.007996132 |
| Uncharacterized protein | C4R0Z8 | 0.002923402 | 0.007970453 | 0.000780232 | 0.003891362 |
| L-type lectin-like domain-containing protein | C4R2L8 | 0.008521138 | 0.00173005 | 0.001235001 | 0.00382873 |
| Metalloprotease with similarity to the zinc carboxypeptidase family | C4R4I2 | 0 | 0.005215868 | 0.002293202 | 0.003754535 |
| Endo-beta-1 3-glucanase major protein of the cell wall involved in cell wall maintenance | C4QYF3 | 0 | 0.003487018 | 0 | 0.003487018 |
| Uncharacterized protein | C4R0V6 | 0.002923402 | 0 | 0 | 0.002923402 |
| 13-beta-glucanosyltransferase | C4QVL4 | 0 | 0.002356878 | 0 | 0.002356878 |
| AP-1 accessory protein | C4R325 | 0 | 0 | 0.002168491 | 0.002168491 |
| Heat shock protein that cooperates with Ydj1p (Hsp40) and Ssa1p (Hsp70) | C4QV89 | 0.003942372 | 0.001490769 | 0.000340725 | 0.001924622 |
| Major exo-13-beta-glucanase of the cell wall involved in cellwall beta-glucan assembly | C4R0Q7 | 0 | 0.001824648 | 0 | 0.001824648 |
| Aminotransferase class I/classII domain-containing protein | C4R862 | 0.003218988 | 0.001592512 | 0.000512162 | 0.001774554 |
| Molecular chaperone | C4QVC4 | 0.001764937 | 0 | 0 | 0.001764937 |
| Superoxide dismutase copper/zinc binding domain-containing protein | C4QW48 | 0.00228556 | 0 | 0.00108123 | 0.001683395 |

### Supplementary Data

| | | | | | |
|---|---|---|---|---|---|
| Protein of the SUN family (Sim1p Uth1p Nca3p Sun4p) that may participate in DNA replication | C4R2Z5 | 0 | 0.001646576 | 0 | 0.001646576 |
| Type II HSP40 co-chaperone that interacts with the HSP70 protein Ssa1p | C4R2Q1 | 0.001411637 | 0.00045896 | 0 | 0.000935299 |
| Sedoheptulose 1 7-bisphosphatase | C4R2M0 | 0.000546974 | 0.000771788 | 0 | 0.000659381 |
| fructose-bisphosphatase | C4R5T8 | 0 | 0.000450901 | 0 | 0.000450901 |
| Xylose and arabinose reductase | C4R135 | 0.000386269 | 0.000487704 | 0 | 0.000436987 |
| Dihydrolipoyl dehydrogenase | C4R312 | 0 | 0.000389153 | 0 | 0.000389153 |
| Serine/threonine-protein phosphatase | C4R1A4 | 0 | 0.000328839 | 0 | 0.000328839 |
| FACT complex subunit | C4QYQ8 | 0.000319168 | 0 | 0 | 0.000319168 |
| Cellulase | C4R8H7 | 0 | 0.000284482 | 0 | 0.000284482 |
| alanine—glyoxylate transaminase | C4R7U0 | 0 | 0.000257802 | 0 | 0.000257802 |
| Transketolase similar to Tkl2p | C4R5P8 | 0.000250801 | 0 | 0 | 0.000250801 |
| Fructose-bisphosphate aldolase | C4QW09 | 0.000246246 | 0 | 0 | 0.000246246 |
| ATPase involved in protein folding and the response to stress. | C4R3X8 | 0 | 0.000214762 | 0 | 0.000214762 |
| Lectin-like protein with similarity to Flo1p thought to be expressed and involved in flocculation | C4QYW7 | 0 | 0.000184489 | 0 | 0.000184489 |
| Plasma membrane Mg(2+) transporter expression and turnover are regulated by Mg(2+) concentration | C4QXF3 | 0 | 0 | 0 | 0 |

#### Supplementary Data

##### Supplementary figures

**Fig. S1.**

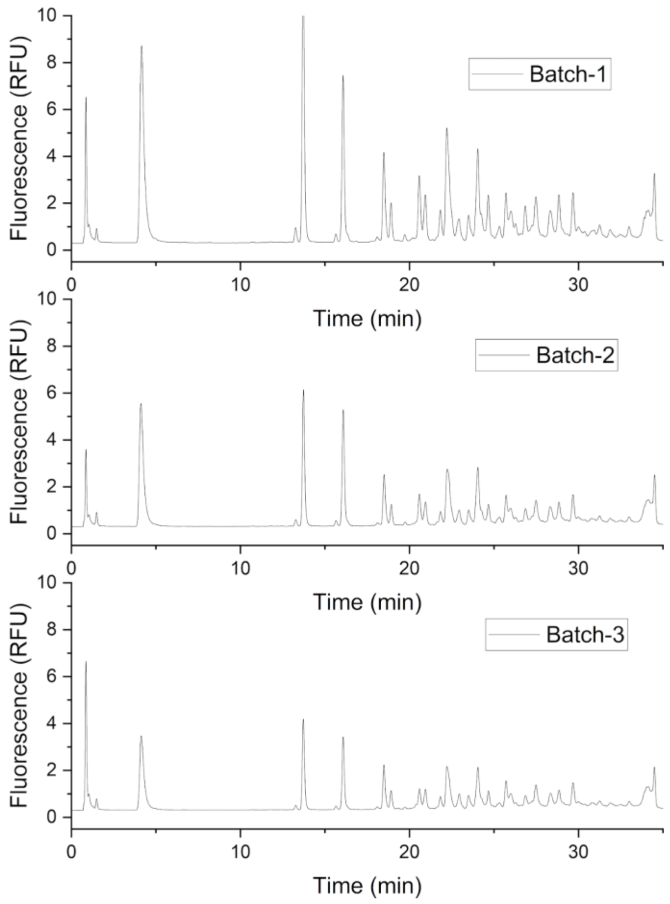


Fig. S1. The HPLC-fluorescence chromatograms of the released N-glycan profiles of three batches of Helaian rhLF. The glycans were released, labeled by InstantPC and measured by UHPLC-Fluoresence detection as described in the Material and Methods section.

**Supplementary Data**

**Fig. S2.**

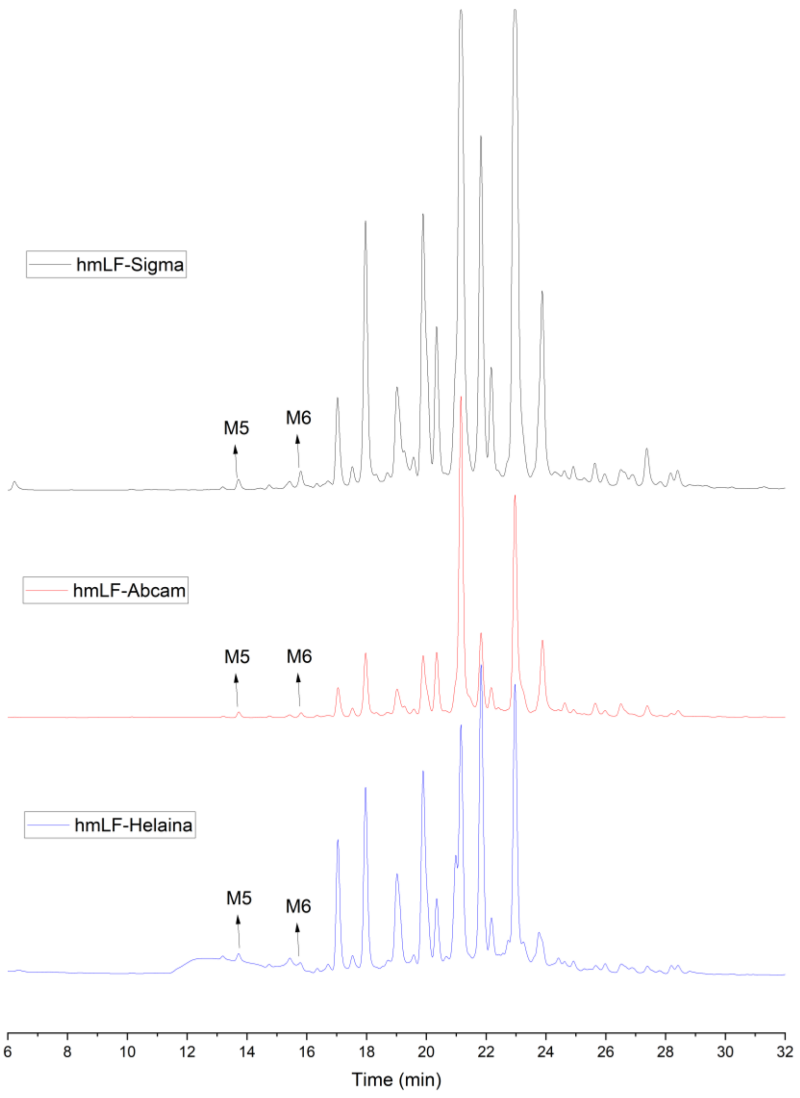


Fig. S2. Comparison of the released N-glycan profiles of native hmLF from three different sources including MilliporeSigma (hmLF-Sigma), Abcam (hmLF-Abcam) and Helaina (hmLF-Helaina). The N-glycans having 5 and 6 mannoses (M5 and M6) were indicated by arrows. The glycans were released, labeled by InstantPC and measured by UHPLC-Fluoresence detection as described in the Material and Methods section.

**Supplementary Data**

**Fig. S3** **(A)**

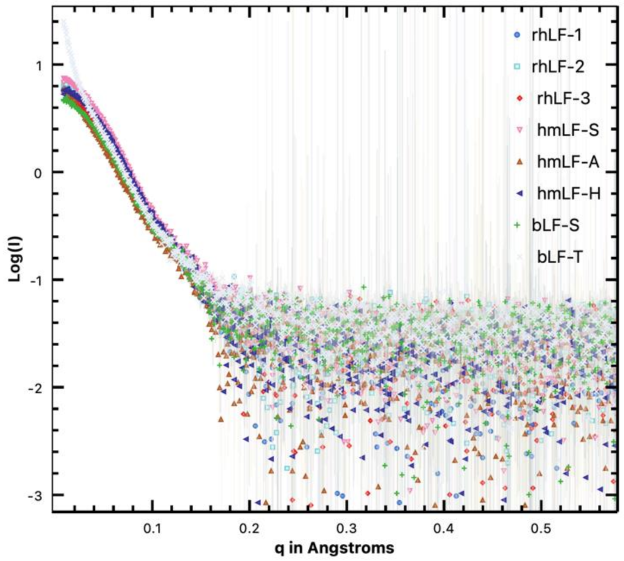


**(B)**

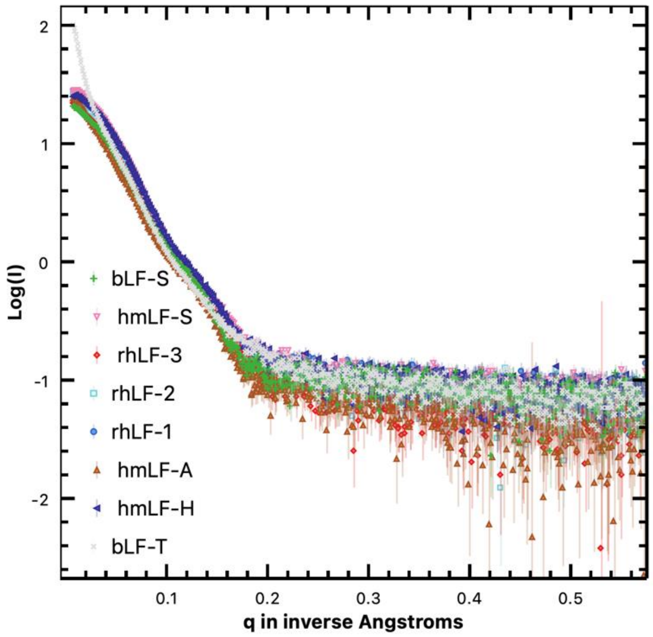


Fig. S3. SAXS raw data of Helaina rhLF (three batches), native hmLF (three sources), and native bLF (two sources) acquired at 1mg/ml (A, top) and 4mg/ml (B, bottom). Helaina rhLF-1, -2, and -3 indicate three batches of Helaina rhLF. hmLF-A, -S, and -H are native hmLF from Abcam, MilliporeSigma, and Helaina, respectively. bLF-S and -T are native bLF supplied by MilliporeSigma and The Lactoferrin Company. Conditions for data acquisition were detailed in the Material and Methods.

**Supplementary Data**

**Fig. S4.**

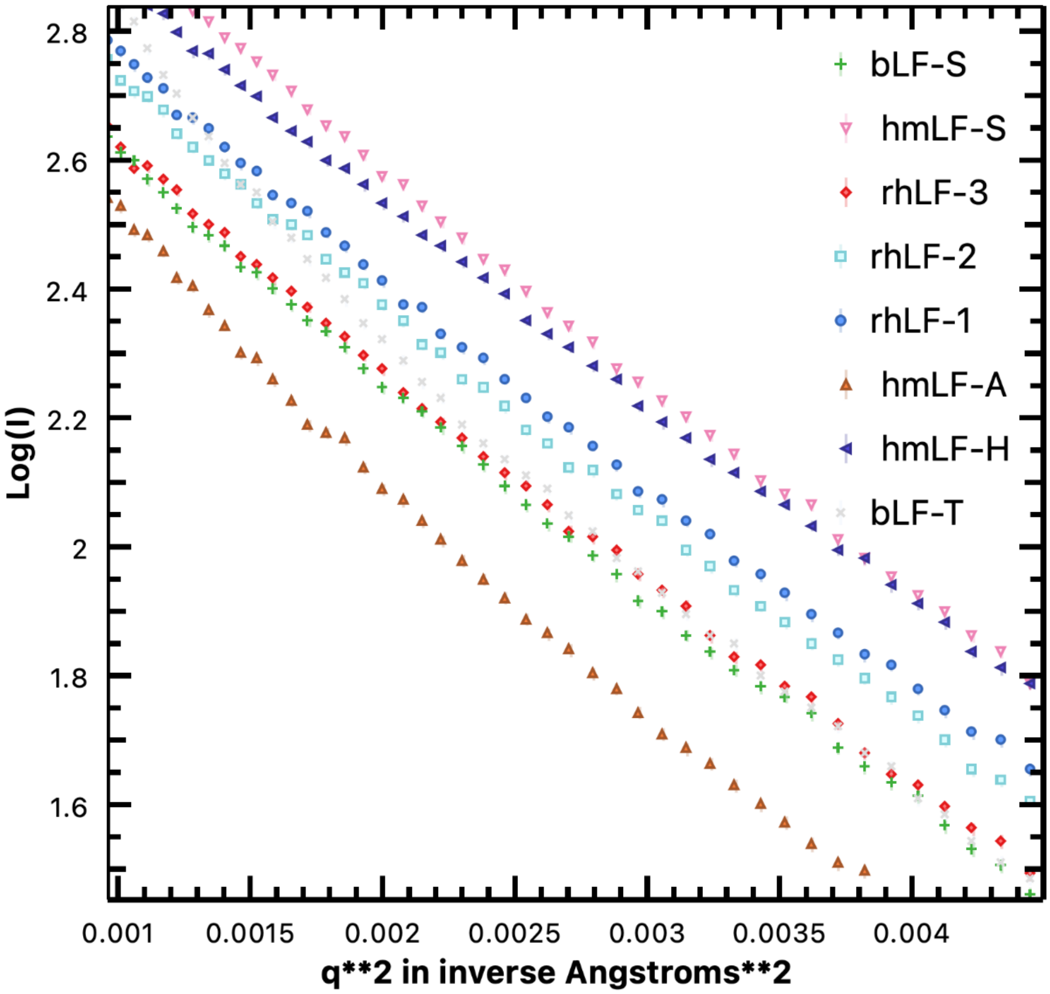


Fig. S4. Guinear plots of Helaina rhLF (three batches), native hmLF (three sources), and native bLF (two sources) derived from the SAXS raw data acquired at 4 mg/mL (Fig. 3S). The sample labels are the same as in Fig. S3. Conditions for data acquisition were detailed in the Material and Methods.

#### Supplementary Data

**Fig. S5.**

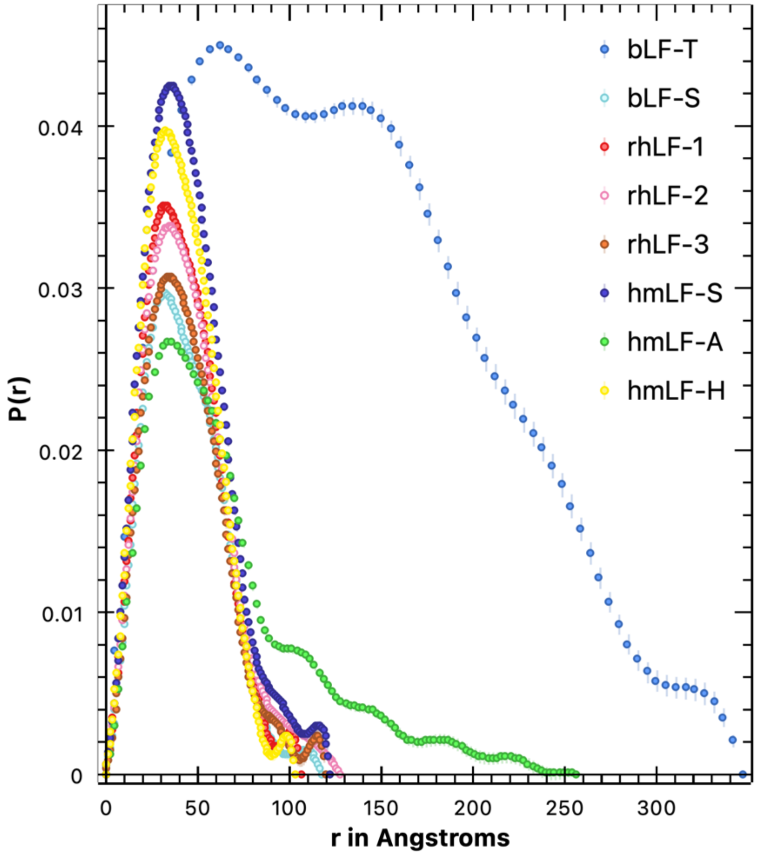


Fig S5, Pair distance distribution function P(r) plot of Helaina rhLF (three batches), native hmLF (three sources), and native bLF (two sources) derived from the SAXS raw data acquired at 4 mg/mL (Fig. 3S). The sample labels are the same as in Fig. S3. Conditions for data acquisition were detailed in the Material and Methods.

**Supplementary Data**

**Fig. S6.**

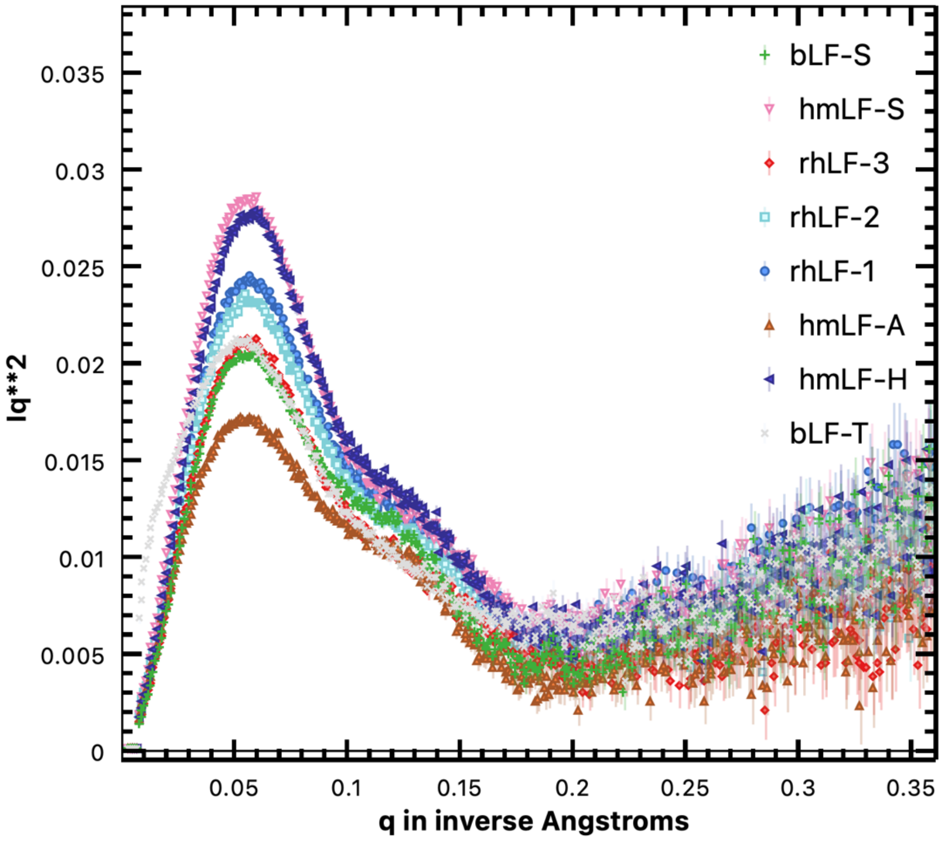


Fig S6, Kratky plots of Helaina rhLF (three batches), native hmLF (three sources), and native bLF (two sources) derived from the SAXS raw data acquired at 4 mg/mL (Fig. 3S). The sample labels are the same as in Fig. S3. Conditions for data acquisition were detailed in the Material and Methods.

#### Supplementary Data

**Fig. S7.**

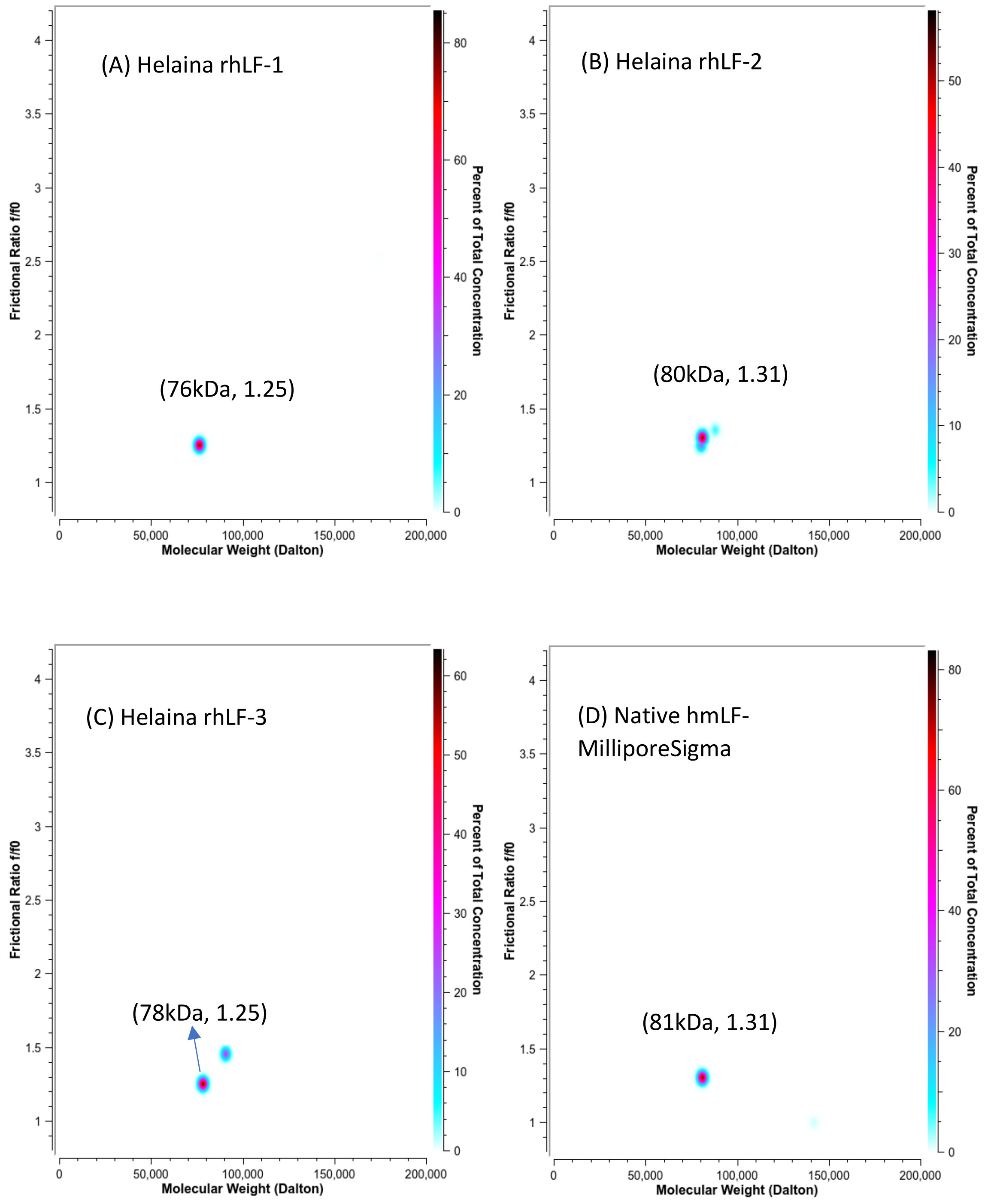

#### Supplementary Data

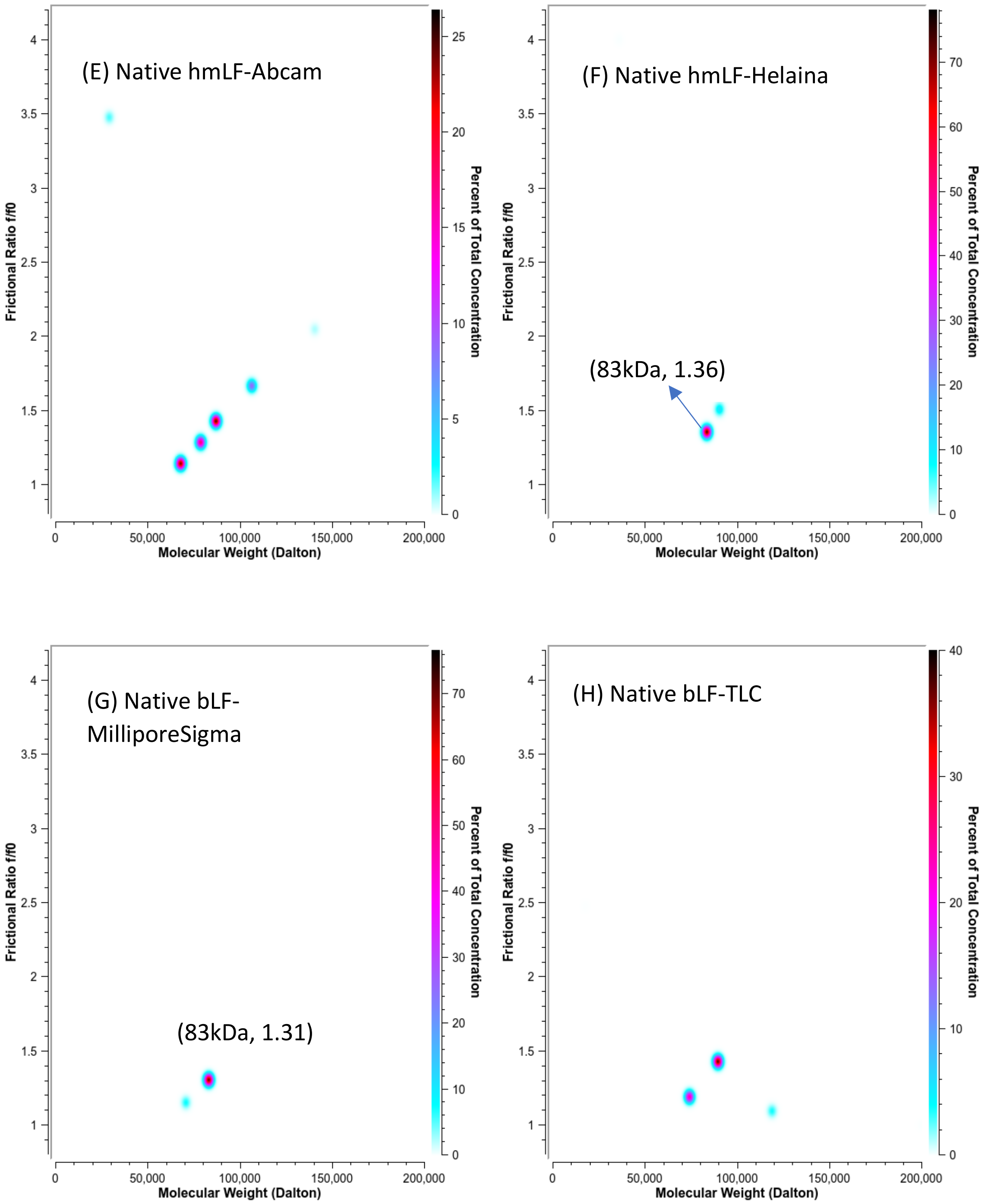


Fig. S7, AUC data of LF protein samples studied at 4 mg/mL concentration. Conditions for data acquisition and processing were detailed in the Material and Methods. The data presented a pseudo-three-dimensional distribution of the observed protein species, with the calculated molecular weight on the x-axis, the calculated frictional ratio on the y-axis, and the % concentration in the Z-plane, with the heat map on the right axis. Conditions for data acquisition and processing were detailed in the Material and Methods.
